# Learning the language of viral evolution and escape

**DOI:** 10.1101/2020.07.08.193946

**Authors:** Brian Hie, Ellen Zhong, Bonnie Berger, Bryan Bryson

## Abstract

Viral mutation that escapes from human immunity remains a major obstacle to antiviral and vaccine development. While anticipating escape could aid rational therapeutic design, the complex rules governing viral escape are challenging to model. Here, we demonstrate an unprecedented ability to predict viral escape by using machine learning algorithms originally developed to model the complexity of human natural language. Our key conceptual advance is that predicting escape requires identifying mutations that preserve viral fitness, or “grammaticality,” and also induce high antigenic change, or “semantic change.” We develop viral language models for influenza hemagglutinin, HIV Env, and SARS-CoV-2 Spike that we use to construct antigenically meaningful semantic landscapes, perform completely unsupervised prediction of escape mutants, and learn structural escape patterns from sequence alone. More profoundly, we lay a promising conceptual bridge between natural language and viral evolution.

**One sentence summary:** Neural language models of semantic change and grammaticality enable unprecedented prediction of viral escape mutations.

## Introduction

Viral mutation that escapes from recognition by neutralizing antibodies has prevented the development of a universal antibody-based vaccine for influenza (Eckert and Kim, 2001; Kim et al., 2018; Krammer, 2019; Kucharski et al., 2015) or human immunodeficiency virus (HIV) (Arrildt et al., 2012; Eckert and Kim, 2001; Richman et al., 2003; Root et al., 2001) and remains an active concern in the development of therapies for COVID-19 (Baum et al., 2020), caused by severe acute respiratory syndrome coronavirus 2 (SARS-CoV-2) infection (Andersen et al., 2020; Walls et al., 2020). Obtaining a better understanding of viral escape has motivated high-throughput experimental techniques, such as deep mutational scans (DMS), that perform causal escape profiling of all single-residue mutations to a viral protein (Dingens et al., 2019; Doud et al., 2018; Lee et al., 2019). Such techniques, however, require substantial effort to profile even a single viral strain, so empirically testing the escape potential of all (combinatorial) mutations in all viral strains remains infeasible.

A more efficient model of viral escape could be achieved computationally. One of our key initial insights is that it may be possible to train an algorithm to learn to model escape from existing viral sequence data alone. Such an approach is not unlike recent algorithmic successes in learning properties of natural language from large text corpuses (Devlin et al., 2018; Peters et al., 2018; Radford et al., 2019); like viral evolution, natural languages like English or Japanese use linear sequence to encode complex concepts (e.g., semantics) and are under complex constraints (e.g., grammar). We pursued the intuition that critical properties of a viral escape mutation have linguistic analogs: first, the mutation must preserve viability and infectivity, i.e., it must be grammatical; second, the mutation must be antigenically altered to evade immunity, i.e., it must have substantial semantic change.

Currently, computational models of viral protein evolution focus on viral fitness (Hopf et al., 2017; Louie et al., 2018) or on functional/antigenic similarity (Alley et al., 2019; Bepler and Berger, 2019; Meroz et al., 2011; Rao et al., 2019) alone. The novel concept critical to our study is that computationally predicting viral escape requires modeling both fitness *and* antigenicity (**Figure 1A**). Moreover, rather than developing two separate models of fitness and function, we reasoned that we could develop a *single* model that *simultaneously* achieves both these tasks. To do so, we leverage state-of-the-art machine learning algorithms (originally developed for natural language understanding) called language models (Devlin et al., 2018; Peters et al., 2018; Radford et al., 2019), which learn the probability of a token (e.g., an English word) given its sequence context (e.g., a sentence) (**Figure 1B**). As done in natural language tasks, we can use a hidden layer output within a neural language model as a semantic embedding (Peters et al., 2018) and the language model output to quantify mutational grammaticality (**Figure 1B**); moreover, the same principles used to train a language model on a sequence of English words can be used to train a language model on a sequence of amino acids.

**Figure 1:**
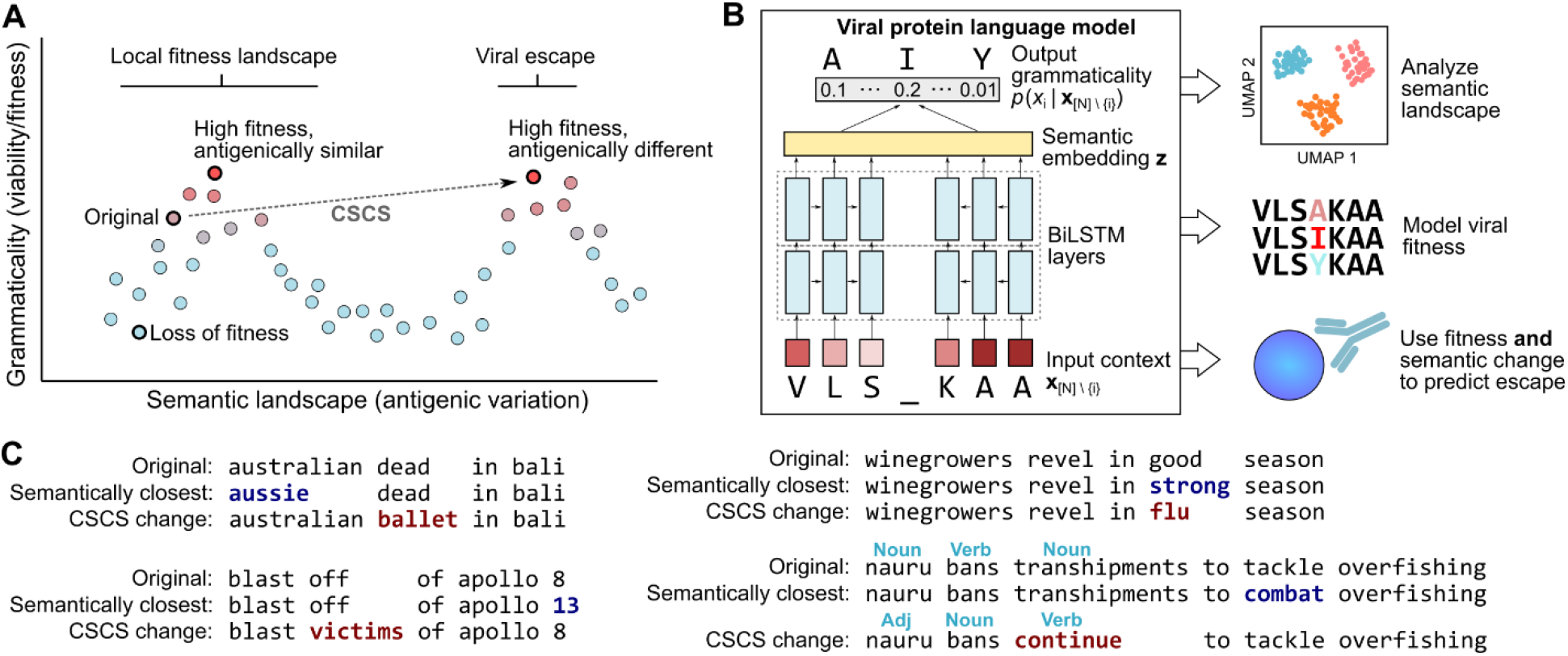
Modeling viral escape requires characterizing semantic change and grammaticality. (**A**) Constrained semantic change search (CSCS) for viral escape prediction is designed to search for mutations to a viral sequence that preserve fitness while being antigenically different. This corresponds to a mutant sequence that is grammatical but has high semantic change with respect to the original (e.g., wildtype) sequence. (**B**) A neural language model with a bidirectional long short-term memory (BiLSTM) architecture is used to learn both semantics (as a hidden layer output) and grammaticality (as the language model output). CSCS combines semantic change and grammaticality to predict escape (**Methods**). (**C**) CSCS-proposed mutations to a news headline (implemented using a neural language model trained on English news headlines) make large changes to the overall semantic meaning of a sentence or to the part-of-speech structure. The semantically closest mutation according to the same model is largely synonymous with the original headline.

Our main hypothesis, therefore, is that (1) language model-encoded semantic change corresponds to antigenic change, (2) language model grammaticality captures viral fitness, and (3) both high semantic change *and* grammaticality help predict viral escape. Searching for mutations with both high grammaticality and high semantic change is a newly formulated task that we call constrained semantic change search (CSCS). For intuitive examples that illustrate the CSCS problem setting, **Figure 1C** contains real CSCS-proposed changes based on a language model trained on news headlines. Additional details of the CSCS problem setup are provided in the **Methods** section.

We propose, to our knowledge, the first general computational model of viral escape. Notably, our neural language model implementation of CSCS is based on sequence data alone (beneficial since sequence is easier to obtain than structure) and requires no explicit escape information (i.e., it is completely unsupervised), does not rely on multiple sequence alignment (MSA) preprocessing (i.e., it is alignment-free), and captures global relationships across an entire sequence (e.g., since word choice at the beginning of a sentence can influence word choice at the end). Additional algorithmic and implementation details are in the **Methods** section.

## Results

Throughout our experiments, we assess generality across viruses by analyzing three important proteins: influenza A hemagglutinin (HA), HIV-1 envelope glycoprotein (Env), and SARS-CoV-2 spike glycoprotein (Spike). All three are found on the viral surface, are responsible for binding host cells, are targeted by antibodies, and are important drug targets given their role in pandemic disease events and widespread human mortality (Andersen et al., 2020; Arrildt et al., 2012; Baum et al., 2020; Eckert and Kim, 2001; Kim et al., 2018; Krammer, 2019; Kucharski et al., 2015; Richman et al., 2003; Root et al., 2001; Walls et al., 2020). We trained a separate language model for each protein using a large corpus of virus-specific amino acid sequences (**Methods**).

We initially sought to investigate the first part of our hypothesis, namely that the semantic embeddings produced by a viral language model would be antigenically meaningful. We computed the semantic embedding for each sequence in the influenza, HIV, and coronavirus corpuses; we then visualized the semantic landscape by learning a two-dimensional approximation of the high-dimensional semantic embedding space using Uniform Manifold Approximation and Projection (UMAP) (McInnes and Healy, 2018). The semantic landscape of each protein shows clear clustering patterns corresponding to subtype, host species, or both (**Figure 2**), suggesting that the model was able to learn functionally meaningful patterns from raw sequence alone.

**Figure 2:**
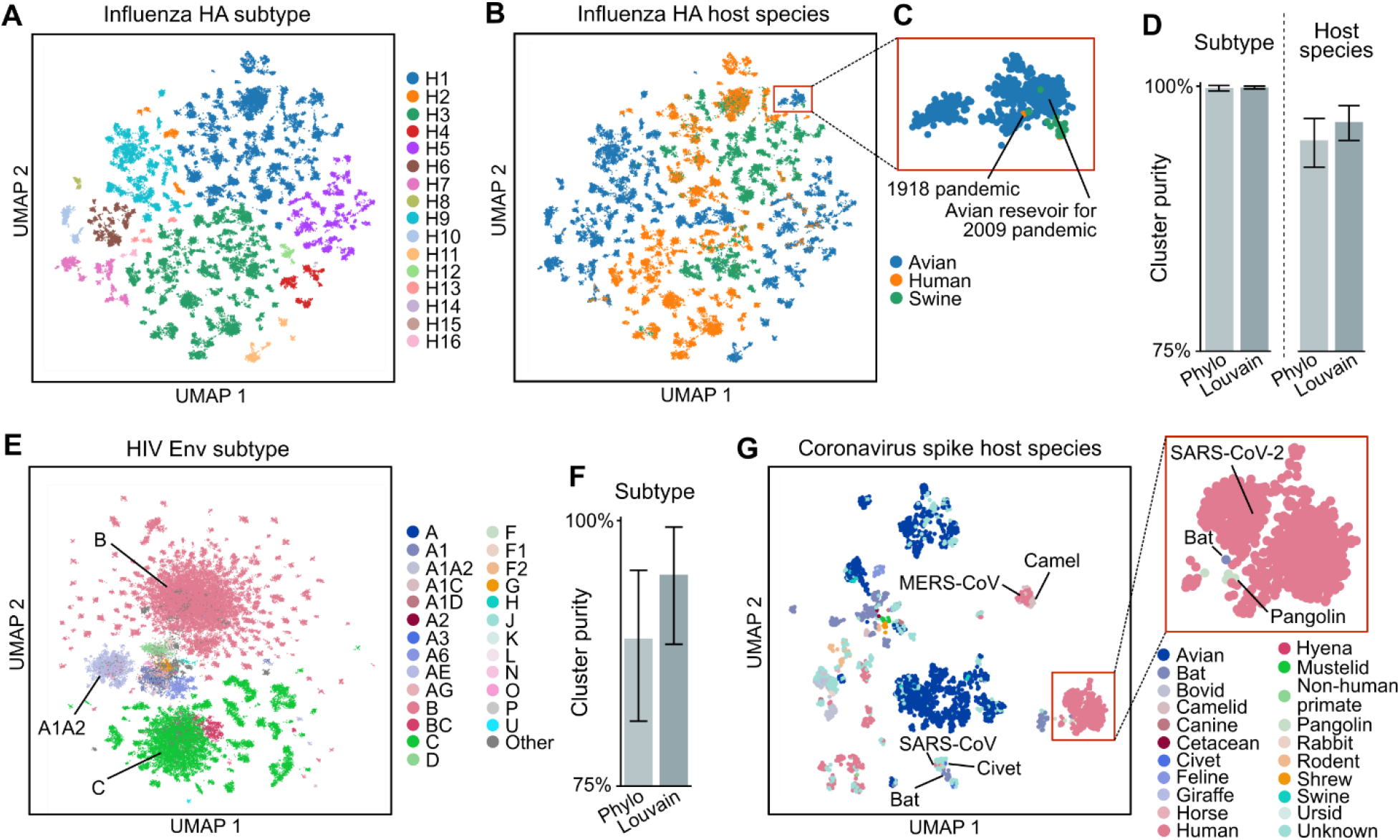
Semantic embedding landscape is antigenically meaningful. (**A, B**) UMAP visualization of the high-dimensional semantic embedding landscape of influenza HA shows clustering structure consistent with subtype and host species. (**C**) A cluster consisting of avian sequences from the 2009 flu season onwards also contains the 1918 pandemic flu sequence, consistent with known antigenic similarity (Wei et al., 2010; Xu et al., 2010). (**D**) Louvain clusters of the HA semantic embeddings have similar purity with respect to subtype or host species compared to a phylogenetic sequence clustering method (Phylo). (**E, F**) The HIV Env semantic landscape shows subtype-related distributional structure, which is supported with high Louvain clustering purity. (**G**) Sequence proximity in the semantic landscape of coronavirus spike proteins is consistent with the possible zoonotic origin of SARS-CoV, MERS-CoV, and SARS-CoV-2.

We can quantify these clear clustering patterns, which are visually enriched for particular subtypes or hosts, by using Louvain clustering (Blondel et al., 2008) to group sequences based on their semantic embeddings (**Figure S1**), followed by measuring the clustering purity based on the percent composition of the most represented metadata category (sequence subtype or host species) within each cluster (**Methods**). Average cluster purities for HA subtype, HA host species, and Env subtype are 99%, 96%, and 95%, respectively, which are comparable to or higher than the clustering purities obtained by more traditional MSA-based phylogenetic reconstruction (Balaban et al., 2019; Katoh and Standley, 2013) (**Figures 2D** and **2F**).

Within the HA landscape, clustering patterns suggest interspecies transmissibility. Interestingly, the sequence for 1918 H1N1 pandemic influenza belongs to the main avian H1 cluster, containing sequences from the avian reservoir for 2009 H1N1 pandemic influenza (**Figures 2C** and **S1**). Our model’s suggested antigenic similarity between H1 HA from 1918 and 2009, though nearly a century apart, has well-established structural and functional support (Wei et al., 2010; Xu et al., 2010). Within the HIV Env landscape, unlike in HA, clusters corresponding to a few subtypes dominate the landscape (**Figure 2E**), perhaps due to the absence of vaccine pressure leading to abundant representation of similar viral strains. Within the landscape of SARS-CoV-2 Spike and homologous proteins, clustering proximity is consistent with the suggested zoonotic origin of several human coronaviruses (**Figure 2G**), including bat and civet for SARS-CoV (Wang and Eaton, 2007); camel for Middle East respiratory syndrome-related coronavirus (MERS-CoV) (Chu et al., 2014); and bat and pangolin for SARS-CoV-2 (Andersen et al., 2020). Analysis of these semantic landscapes strengthens our hypothesis that our viral sequence embeddings encode functional and antigenic variation.

Not only does escape prediction stand to benefit from modeling antigenic change, but from modeling viral fitness as well. Therefore, in line with the second part of our hypothesis, we assessed the relationship between viral fitness and language model grammaticality using high-throughput DMS characterization of hundreds or thousands of mutants to a given viral protein. We obtained datasets measuring replication fitness of all single-residue mutations to A/WSN/1933 (WSN33) HA H1 (Doud and Bloom, 2016), combinatorial mutations to antigenic site B in six HA H3 strains (Wu et al., 2020), or all single-residue mutations to BG505 and BF520 HIV Env (Haddox et al., 2018), as well as a dataset measuring the dissociation constant (Kd) between combinatorial mutations to SARS-CoV-2 Spike and human ACE2 (Starr et al., 2020), which we use to approximate the fitness of Spike.

We found that language model grammaticality was significantly correlated with viral fitness consistently across all viral strains and across studies that performed single or combinatorial mutations (**Figure 3A** and **Table S1**), even though our language models were not given any explicit fitness-related information and were not trained on the DMS mutants. Strikingly, when we instead compared viral fitness to the magnitude of mutant semantic change (rather than grammaticality), we observed significant *negative* correlation in nine out of ten strains tested (**Figure 3A** and **Table S1**). This makes sense biologically, since a mutation with a large effect on function is on average more likely to be deleterious and result in a loss of fitness. These results suggest that, as hypothesized, grammatical “validity” of a given mutation captures fitness information, and adds an additional dimension to our understanding of how semantic change encodes perturbed protein function.

**Figure 3:**
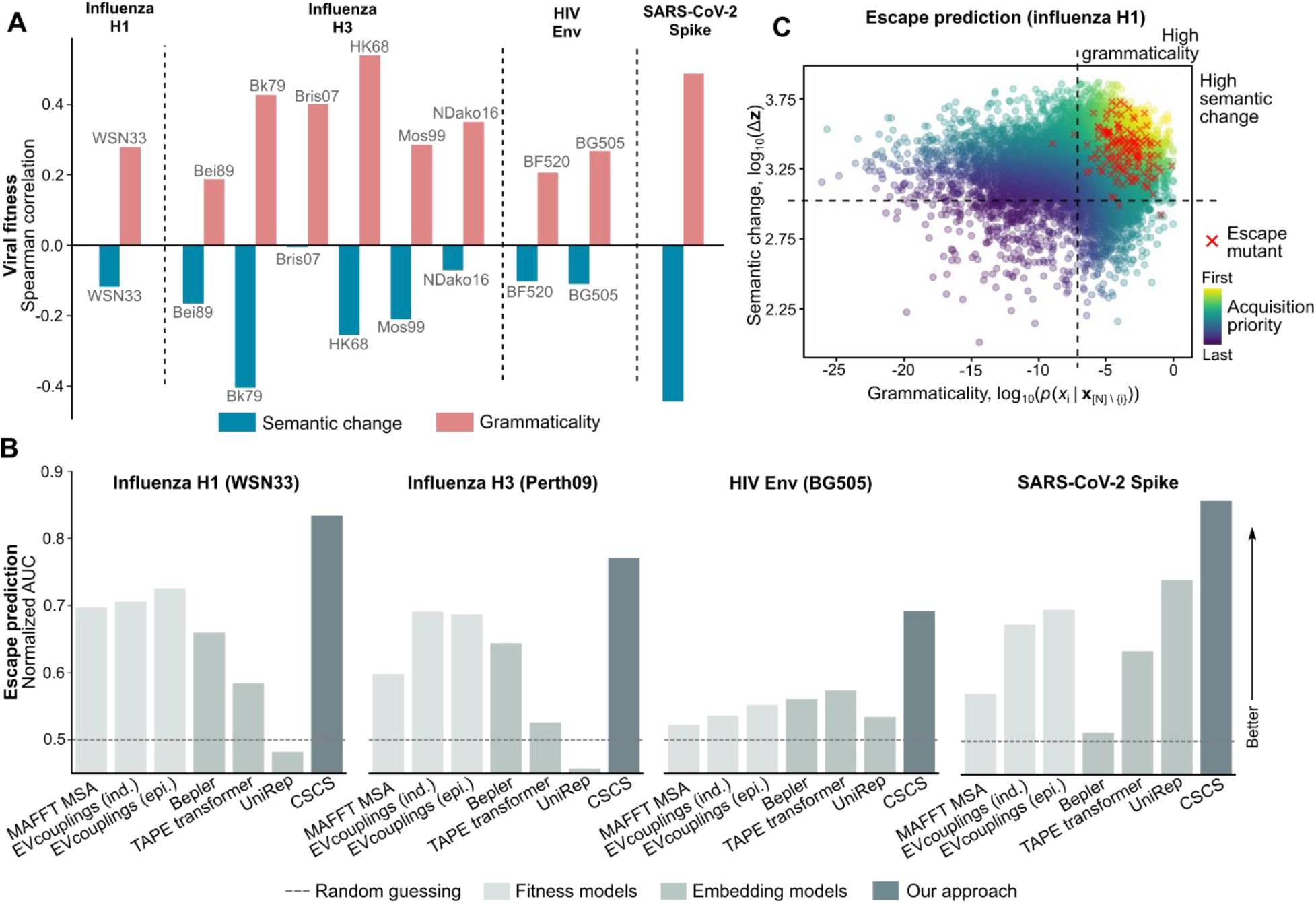
Biological interpretation of language model semantics and grammaticality enables escape prediction. (**A**) While grammaticality is positively correlated with the results of DMS viral fitness measurements, semantic change is negatively correlated with fitness, suggesting that most semantically altered proteins lose fitness. Influenza strains are A/WSN/1933 (WSN33), A/Hong Kong/1/1968 (HK68), A/Bangkok/1/1979 (Bk79), A/Beijing/353/1989 (Bei89), A/Moscow/10/1999 (Mos99), A/Brisbane/10/2007 (Bris07), and A/North Dakota/26/2016 (NDako16). (**B, C**) However, when a mutation is ensured to have both high semantic change *and* high grammaticality, it is more likely to induce escape. Considering both semantic change and grammaticality enables enriched acquisition of escape mutants that is consistently higher than that of previous fitness models or generic functional embedding models.

Based on these promising analyses of viral semantics and grammaticality, we therefore sought to test the third part of our hypothesis, namely that combining semantic change and grammaticality enables escape mutation prediction. Our experimental setup initially involves making, *in silico*, all possible single-residue mutations to a given viral protein sequence; then, each mutant is ranked according to the CSCS objective that combines semantic change and grammaticality. We validate this ranking based on enriched CSCS acquisition of experimentally verified mutants that causally induce escape from neutralizing antibodies. Three of these causal escape datasets used DMS followed by antibody selection to identify escape mutants to WSN33 HA H1 (Doud et al., 2018), A/Perth/16/2009 (Perth09) HA H3 (Lee et al., 2019), and BG505 Env (Dingens et al., 2019). The fourth identified escape mutants to Spike based on natural replication error after two *in vitro* passages under antibody selection (Baum et al., 2020), in contrast to a more exhaustive DMS.

To quantify enrichment of CSCS-acquired escapes, we compute the area under the curve (AUC) of the number of acquired escape mutations versus the total acquired mutations, normalized to be between 0 and 1, where a value of 0.5 indicates random guessing and higher values indicate greater enrichment. In all four cases, escape prediction with CSCS results in both statistically significant and strong AUCs of 0.834, 0.771, 0.692, and 0.856 for H1 WSN33, H3 Perth09, Env BG505, and Spike, respectively (one-sided permutation-based *P* < 1 × 10^−5^ in all cases) (**Figure 3B** and **Table S2**). We emphasize that none of the escape mutants are present in the training data, and we did not provide the model with any explicit information on escape, a challenging problem setup in machine learning referred to as “zero-shot prediction” (Radford et al., 2019).

Crucially, in support of our hypothesis, the escape AUC strictly decreases when ignoring either grammaticality or semantic change, evidence that *both* are useful in predicting escape (**Figures 3C** and **S2**, **Table S2**). Note that while semantic change is negatively correlated with fitness, it is positively predictive (along with grammaticality) of escape (**Table S2**); the analogous biological interpretation is that functional mutations are often deleterious but, when fitness is preserved, they are associated with antigenic change and subsequent escape from immunity.

For a benchmark comparison, we also tested how well alternative models of fitness (each requiring MSA preprocessing) or of semantic change (pretrained on generic protein sequence) predict escape, noting that these models were not explicitly designed for escape prediction. We found that CSCS with our viral language models was substantially more predictive of causal escape mutants in all four viral proteins (**Figure 3B**). Moreover, the individual grammaticality or semantic change components of our language models often outperformed the corresponding benchmark models (**Table S2**), demonstrating the value of nonlinear, high-capacity fitness models or of virus-specific, finetuned semantic embedding models, respectively. In total, our results provide strong empirical support for the hypothesis that both semantic change and grammaticality are useful for escape prediction.

A notable aspect of our results is that, though viral escape is mechanistically linked to a viral protein’s structure (Dingens et al., 2019; Doud et al., 2018; Lee et al., 2019), our models are trained entirely from sequence and bypass explicit structural information altogether (which is often difficult to obtain). Given our validated escape prediction capabilities, we wanted to look at our model’s escape predictions in the context of three-dimensional protein structure to see if our model was able to learn structurally relevant patterns from sequence alone. We used CSCS to score each residue based on predicted escape potential, from which we could visualize escape potential across the protein structure and quantify significant enrichment or depletion of escape potential (**Methods**).

For both HA H1 and H3, we found that escape potential is significantly enriched in the HA head and significantly depleted in the HA stalk (**Figures 4A, 4B**, and **S3**; **Table S3**), consistent with existing literature on HA mutation rates and supported by the successful development of anti-stalk broadly neutralizing antibodies (Ekiert et al., 2009; Kallewaard et al., 2016). Also consistent with existing knowledge is the significant enrichment of escape mutations in the V1/V2 hypervariable regions of Env (**Figures 4C, 4D**, and **S3**; **Table S3**) (Sagar et al., 2006); interestingly, our model also associates significant antigenic change with the gp120 inner domain. An important point is that our model only learns escape patterns that can be linked to mutations, rather than post-translational changes like glycosylation that contribute to HIV escape (Wei et al., 2003), which may explain the lack of statistically significant escape potential assigned to Env glycosylation sites (**Figure 4C** and **Table S3**).

**Figure 4:**
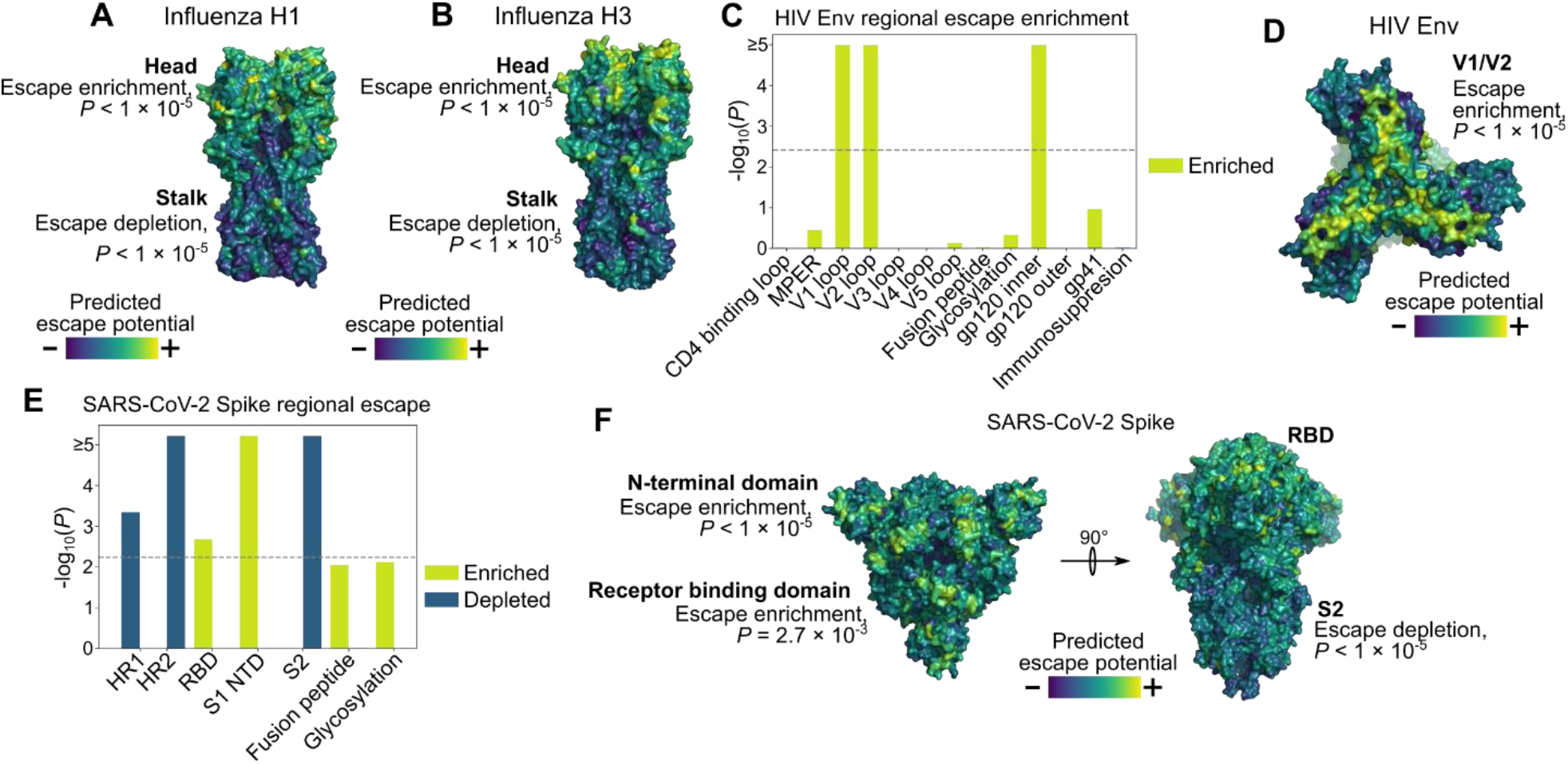
Structural localization of predicted escape potential. (**A, B**) HA trimer colored by escape potential. Escape potential is significantly enriched in the HA head but significantly depleted in the HA stalk. (**C**) Escape potential in HIV Env is significantly enriched in the V1 and V2 hypervariable regions and in the inner domain of gp120; gray dashed line indicates statistical significance threshold. (**D**) The Env trimer colored by escape potential, oriented to show the V1/V2 regions. (**E**) Potential for escape in SARS-CoV-2 Spike is significantly enriched at the N-terminal domain and receptor binding domain (RBD) and significantly depleted at multiple regions in the S2 subunit; gray dashed line indicates statistical significance threshold. (**F**) The Spike trimer colored by escape potential, oriented to show the RBD (left) and S2 (right).

Interestingly, the escape potential within SARS-CoV-2 Spike is significantly enriched in both the receptor binding domain (RBD) and N-terminal domain, while escape potential is significantly depleted in the S2 subunit (**Figures 4E, 4F**, and **S3**; **Table S3**), results which are further supported by greater evolutionary conservation at S2 antigenic sites (Ravichandran et al., 2020). Our model of Spike escape therefore suggests that immunodominant antigenic sites in S2 (Baum et al., 2020; Ravichandran et al., 2020) may be more stable target epitopes for antibody-based neutralization and underscores the need for more exhaustive causal escape profiling of Spike, especially in regions aside from the RBD.

Predicted escape potential is therefore highly specific to particular regions of protein structure even when (as in the HA stalk domain) these structures correspond to non-contiguous regions of protein sequence. Our language models therefore learn structurally-relevant patterns, as well as information on how this structure interacts with its environment, from sequence alone.

## Discussion

Our study offers strong evidence that viral language models provide a new and rich analytic strategy to be used alongside traditional tools based on sequence alignment and phylogeny. Continuous embeddings of antigenic semantics could have a number of downstream applications, such as leveraging diversity-preserving subsampling techniques (Hie et al., 2019) to select components of a multivalent or mosaic vaccine (Barouch et al., 2018; Boyoglu-Barnum et al., 2020). CSCS could also be applied to other problems, such as drug resistance, in which a combination of grammaticality and semantic change would be valuable. A more profound implication is that the “distributional hypothesis” from linguistics (Harris, 1954), in which co-occurrence patterns can model complex concepts and on which language models are based, can be extended to viral evolution, establishing a hopefully productive dialogue between two disparate fields.

## Acknowledgements

We thank Alejandro Balazs, Owen Leddy, Adam Lerer, Allen Lin, Adam Nitido, Uma Roy, and Aaron Schmidt for helpful discussions. We thank Steven Chun, Benjamin DeMeo, Ashwin Narayan, An Nguyen, Sarah Nyquist, and Alexander Wu for assistance with the manuscript.

## Funding

B.H. and E.Z. are partially supported by NIH grant R01 GM081871 (to B. Berger). B.H. is partially supported by the Department of Defense (DoD) through the National Defense Science and Engineering Graduate Fellowship (NDSEG). E.Z. is partially supported by the National Science Foundation (NSF) Graduate Research Fellowship.

## Author contributions

All authors conceived the project and methodology. B.H. performed the computational experiments and wrote the software. All authors interpreted the results and wrote the manuscript.

## Competing interests

The authors declare no competing interests.

## Data and code availability

Code, data, and pretrained models are available at https://github.com/brianhie/viral-mutation. We used the following publicly available datasets for model training:

- Influenza A HA protein sequences from the NIAID Influenza Research Database (IRD) (http://www.fludb.org)
- HIV-1 Env protein sequences from the Los Alamos National Laboratory (LANL) HIV database (https://www.hiv.lanl.gov)
- *Coronavidae* spike protein sequences from the Virus Pathogen Resource (ViPR) database (https://www.viprbrc.org/brc/home.spg?decorator=corona)
- SARS-CoV-2 Spike protein sequences from NCBI Virus (https://www.ncbi.nlm.nih.gov/labs/virus/vssi/)
- SARS-CoV-2 Spike and other Betacoronavirus spike protein sequences from GISAID (https://www.gisaid.org/)

We used the following publicly available datasets for fitness and escape validation:

- Fitness single-residue DMS of HA H1 WSN33 from Doud and Bloom (2016) (Doud and Bloom, 2016)
- Fitness combinatorial DMS of antigenic site B in six HA H3 strains from Wu et al. (Wu et al., 2020)
- Fitness single-residue DMS of Env BF520 and BG505 from Haddox et al. (Haddox et al., 2018)
- ACE2 binding affinity combinatorial DMS of SARS-CoV-2 from Starr et al. (Starr et al., 2020)
- Escape single-residue DMS of HA H1 WSN33 from Doud et al. (2018) (Doud et al., 2018)
- Escape single-residue DMS of HA H3 Perth09 from Lee et al. (Lee et al., 2019)
- Escape single-residue DMS of Env BG505 from Dingens et al. (Dingens et al., 2019)
- Escape mutations of Spike from Baum et al. (Baum et al., 2020)

Links to processed training and validation datasets are also available at https://github.com/brianhie/viral-mutation.

## Methods

### CSCS: Problem Formulation

Intuitively, our goal is to identify mutations that induce high semantic change (e.g., a large impact on biological function) while being grammatically acceptable (e.g, biologically viable). More precisely, we are given a sequence of tokens defined as 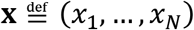 such that 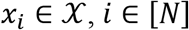, where 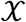 is a finite alphabet. Let 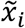 denote a mutation at position *i* and the mutated sequence as 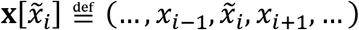.

We first require a semantic embedding 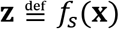, where 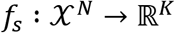 embeds discrete-alphabet sequences into a *K*-dimensional continuous space, where closeness in embedding space corresponds to semantic similarity. We denote semantic change as the distance in embedding space, i.e.,

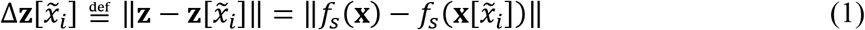

where ‖∙‖ denotes a vector norm. The grammaticality of a mutation is described by

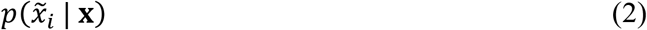

which takes values close to zero if 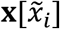 is not grammatical and close to one if it is grammatical.

Our objective combines semantic change and grammaticality. Taking inspiration from upper confidence bound acquisition functions in Bayesian optimization (Auer, 2003), which additively weigh a predicted value with its uncertainty, we can combine terms *(1)* and (2) with a weight parameter *β* ∈ [0, ∞) above to compute

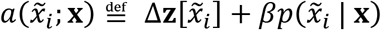

for each possible mutation 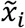. 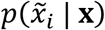 can be interpreted as the “uncertainty” associated with a given semantic shift 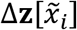. Mutations 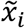 are prioritized based on 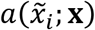; we refer to this ranking of mutations based on semantic change and grammaticality as CSCS.

### CSCS: Algorithms

Algorithms for CSCS could potentially take many forms; for example, separate algorithms could be used to compute 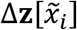 and 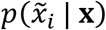 independently, or a two-step approach might be possible that computes one of the terms based on the value of the other.

Instead, we reasoned that a single approach could compute both terms simultaneously, based on learned language models that learn the probability distribution of a word given its context (Dai and Le, 2015; Devlin et al., 2018; Mikolov et al., 2013; Peters et al., 2018; Radford et al., 2019). The language model we use throughout our experiments considers the full sequence context of a word and learns a latent variable probability distribution 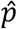 and function 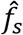 over all *i* ∈ [*N*] where

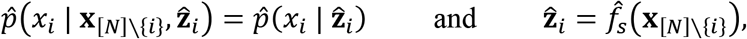

i.e., latent variable 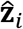 encodes the sequence context 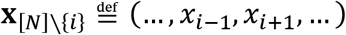 such that *x*_*i*_ is conditionally independent of its context given the value of 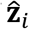.

We use different aspects of the language model to describe semantic change and grammaticality by setting terms *(1)* and (2) as

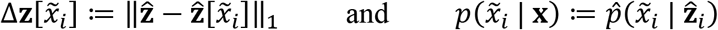

Where 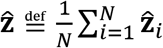 is the average embedding across all positions, 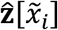 is defined similarly but for the mutated sequence, and ‖∙‖_1_ is the *ℓ*_1_ norm. Effectively, distances in embedding space encode semantic change and the emitted probability encodes grammaticality.

Based on the success of recurrent architectures for protein-sequence representation learning (Alley et al., 2019; Bepler and Berger, 2019; Rao et al., 2019), we use similar encoder models for viral protein sequences (**Figure 1B**). Our model passes the full context sequence into BiLSTM hidden layers. Null character pre-padding was used to handle variable-length sequences. We used the concatenated output of the final LSTM layers as the semantic embedding, i.e.,

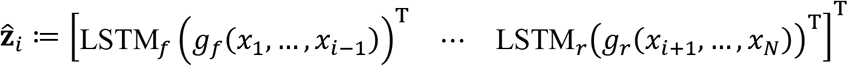

Where *g_f_* is the output of the preceding forward-directed layer, LSTM_*f*_ is the final forward-directed LSTM layer, and *g*_*r*_ and LSTM_*r*_ are the corresponding reverse-directed components. The final output probability is a softmax-transformed linear transformation of 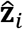, i.e.,

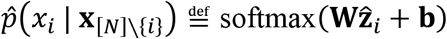

for some learned model parameters **W** and **b**. In our experiments, we used a 20-dimensional learned dense embedding for each element in the alphabet 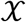, two BiLSTM layers with 512 units, and categorical cross entropy loss optimized by Adam with a learning rate of 0.001, *β*_1_ = 0.9, and *β*_2_ = 0.999. Hyperparameters and architecture were selected based on a small-scale grid search as described below.

Rather than acquiring mutations based on raw semantic change and grammaticality values, which may be on very different scales, we find that calibrating *β* is much easier in practice when first rank-transforming the semantic change and grammaticality terms, i.e., acquiring based on

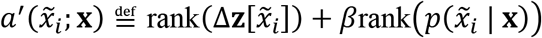

All possible mutations 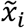 are then given priority based on the corresponding values of 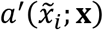, from highest to lowest. Our empirical results seem consistently well-calibrated around *β* = 1 (equally weighting both terms), which we used in all of our experiments.

### CSCS: Extension to combinatorial mutations

The above exposition is limited to the setting in which mutations are assumed to be single-token. We perform a simple extension to handle combinatorial mutations. We denote such a mutant sequence as 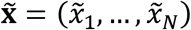, which has the same length as **x**, where the set of mutations consists of the tokens in 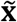 that disagree with those at the same position in **x**, which we denote

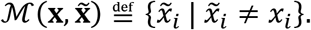

The semantic embedding can simply be computed as 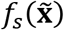 from which semantic change can be computed as above. For the grammaticality score, we make a simple modeling assumption and compute grammaticality as

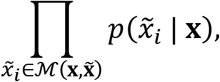

i.e., the product of the probabilities of the individual point-mutations. We note that this works well empirically in the combinatorial fitness datasets that we test, even when the number of mutations is not fixed as in the SARS-CoV-2 DMS Kd dataset. Other ways of estimating joint, combinatorial grammaticality terms while preserving efficient inference are also worth considering in future work.

While we do not consider insertions or deletions in this study, we do note that, in viral sequences, insertions and deletions are rarer than substitutions by a factor of four or more (Sanjuán et al., 2010) and the viral mutation datasets that we considered exclusively profiled substitution mutations alone. Extending our algorithms to compute semantic change of sequences with insertions or deletions would be essentially unchanged from above. The more difficult task is in reasoning about and modeling the grammaticality of an insertion or a deletion. While various grammaticality heuristics based on the language model output may be possible, this is also an interesting area for further methodological development.

### Language model selection and training details

To select the model architecture described above, we performed a small-scale grid search using categorical cross entropy loss on the influenza dataset described below. We evaluated language model performance with a test set of held-out HA sequences where the first recorded date was before 1990 or after 2017, yielding a test set of 7,497 out of 44,999 sequences (about 17%). Hyperparameter search ranges were influenced by previous applications of recurrent architectures to protein sequence representation learning (Bepler and Berger, 2019). We tested hidden unit dimensions of 128, 256, and 512. We tested architectures with one or two hidden layers. We tested three hidden-layer architectures: a densely connected neural network with access to both left and right sequence contexts, an LSTM with access to only the left context, and a BiLSTM with access to both left and right sequence contexts. We tested two Adam learning rates (0.01 and 0.001). All other architecture details were fixed to reasonable defaults. In total, we tested 36 conditions, and ultimately used a BiLSTM architecture with two hidden layers of 512 hidden units each, with an Adam learning rate of 0.001. We used the same architecture for all experiments. We train the language model to predict the observed amino acid residue at all positions in each sequence, using the remaining sequence as the input; one training epoch is completed when the model has considered all positions in all sequences in the training corpus. We trained each model until convergence of cross entropy loss across one training epoch.

### News headline data, model training, and CSCS

Preprocessed headlines (stripped of punctuation, space-delimited, and lower-cased) from the Australian Broadcasting Corporation (early-2013 through the end of 2019) were obtained from https://www.kaggle.com/therohk/million-headlines. We trained a word-level BiLSTM language model. Semantic embeddings and grammaticality were quantified as described above. To obtain CSCS-proposed headline mutations, we considered all possible single-word mutations and acquired the top according to the CSCS objective. For comparison, we also acquired the single-word mutated headline with the closest embedding vector to the original headline as the “semantically closest” mutation.

### Viral protein sequence datasets and model training

We trained three separate language models for influenza HA, HIV Env, and SARS-CoV-2 Spike using the model architecture described above. One training epoch consisted of predicting each token over all sequences in the training set.

Influenza HA amino acid sequences were downloaded from the “Protein Sequence Search’” section of https://www.fludb.org. We only considered complete hemagglutinin sequences from virus type A. We trained an amino acid residue-level language model on a total of 44,851 unique influenza A hemagglutinin (HA) amino acid sequences observed in animal hosts from 1908 through 2019.

HIV Env protein sequences were downloaded from the “Sequence Search Interface’” at the Los Alamos National Laboratory (LANL) HIV database (https://www.hiv.lanl.gov) (Foley et al., 2018). All complete HIV-1 Env sequences were downloaded from the database, excluding sequences that the database had labeled as “problematic.” We additionally only considered sequences that had length between 800 and 900 amino acid residues, inclusive. We trained an amino acid residue-level language model on a total of 57,730 unique Env sequences.

*Coronavidae* spike glycoprotein sequences were obtained from the Gene/Protein Search portal of the ViPR database (https://www.viprbrc.org/brc/home.spg?decorator=corona) across the entire *Coronavidae* family. We only included amino acid sequences with “spike” gene products. SARS-CoV-2 Spike sequences were obtained from the Severe acute respiratory syndrome coronavirus 2 datahub at NCBI Virus (https://www.ncbi.nlm.nih.gov/labs/virus/vssi/). Betacoronavirus spike sequences from GISAID also used in Starr et al.’s analysis (Starr et al., 2020) were obtained from https://github.com/jbloomlab/SARS-CoV-2-RBD_DMS/blob/master/data/alignments/Spike_GISAID/spike_GISAID_aligned.fasta. Across all coronavirus datasets, we furthermore excluded sequences with a protein sequence length of less than 1,000 amino acid residues. We trained an amino acid residue-level language model on a total of 4,172 unique Spike (and homologous protein) sequences.

### Semantic embedding landscape visualization, clustering, and quantification

We used the language models for HA, Env, and Spike to produce semantic embeddings for sequences within each language model’s respective training corpus, where the semantic embedding procedure is described above. We used the UMAP (McInnes and Healy, 2018) Python implementation (https://github.com/lmcinnes/umap) as wrapped by the Scanpy version 1.4.5 Python package (Wolf et al., 2018) (https://scanpy.readthedocs.io/) using the 100-nearest neighbors graph for influenza and HIV and the 20-nearest neighbors graph for coronavirus and with a minimum distance parameter of 1 for all embeddings.

We used Louvain clustering (Blondel et al., 2008) with a resolution parameter of 1, also using the implementation wrapped by Scanpy, to cluster sequences within each viral corpus based on the same nearest neighbors graph used to compute the UMAP embedding. Louvain cluster purity was evaluated with respect to a metadata class (e.g., host species or subtype) by first calculating the percent composition of each metadata class label within a given cluster and using the maximum composition over all class labels as the purity percentage; we calculated this purity percentage for each Louvain cluster.

To compare Louvain clustering purities to a more traditional sequence clustering strategy, we used MAFFT version 7.453 (Katoh and Standley, 2013) (https://mafft.cbrc.jp/alignment/software/) to construct a phylogenetic tree of the respective viral sequence corpus. We then use TreeCluster (Balaban et al., 2019) (https://github.com/niemasd/TreeCluster) to group sequences based on the MAFFT tree. We performed a range search of the TreeCluster threshold parameter to ensure that the number of returned clusters was equal to the number of clusters returned by the Louvain clustering of the same sequence corpus; all other parameters were set to the defaults. Using the cluster labels returned by TreeCluster, we computed cluster purities using the same procedure described for Louvain clustering above.

### Fitness validation

We obtained mutational fitness preference scores for HA H1 WSN33 mutants from Doud and Bloom (Doud and Bloom, 2016), preference scores for antigenic site B mutants in six HA H3 strains (Bei89, Bk79, Bris07, HK68, Mos99, NDako16) from Wu et al. (Wu et al., 2020), preference scores for Env BF520 and BG505 mutants from Haddox et al. (Haddox et al., 2018), and Kd binding affinities between SARS-CoV-2 mutants and ACE2 from Starr et al. (Starr et al., 2020). For the replication fitness DMS data, we used the preference scores averaged across technical replicates (as done within each study). For the ACE2 DMS data, if a sequence had more than one measured Kd, we took the median Kd as the representative fitness score for that sequence.

Using the corresponding pretrained viral fitness model, we computed the grammaticality score and semantic change (with respect to the original wildtype sequence) for each mutant sequence produced in each of the above studies. We computed the Spearman correlation between fitness preference or binding Kd with either the grammaticality score or the semantic change. We used the implementation provided in the scipy version 1.3.1 Python package (https://www.scipy.org/) to compute Spearman correlation coefficients and corresponding *P* values.

### Escape prediction validation

We obtained experimentally validated causal escape mutants to HA H1 WSN33 from Doud et al. (Doud et al., 2018), HA H3 Perth09 from Lee et al. (Lee et al., 2019), Env BG505 from Dingens et al. (Dingens et al., 2019), and Spike from Baum et al. (Baum et al., 2020). We then made, *in silico*, all possible single-residue mutations to H1 WSN33, H3 Perth09, Env BG505, and Spike. For each of these mutations, we computed semantic change and grammaticality and combined these scores using the CSCS rank-based acquisition function. For a given viral protein, the value of the CSCS acquisition function was used to rank all possible mutants. To assess enrichment of acquired escape mutants, we constructed a curve that plotted the top *n* CSCS-acquired mutants on the x-axis and the corresponding number of these mutants that were also causal escape mutations on the y-axis; the area under this curve, normalized to the total possible area, resulted in our normalized AUC metric for evaluating escape enrichment.

We computed a permutation-based *P* value to assess the statistical significance of the enrichment of a given CSCS ranking. To do so, we constructed a null distribution by randomly sampling (without replacement) a subset of mutants as a “null escape” set, controlling for the number of mutants by ensuring that the null escape mutant set was the same size as the true escape mutant set, and recalculating the normalized AUC accordingly (essentially “permuting” the escape versus non-escape labels). We repeated this for 100,000 permutations. Bonferroni-corrected *P* values were considered statistically significant if they were below 0.05.

### Escape prediction benchmarking

We wanted to benchmark our ability to predict escape, which is based on combining grammaticality and semantic change, based on previous methods that assess either viral fitness (which we conceptually link to grammaticality) or learn functional representations (which we conceptually link to semantics) alone.

For our first fitness model, we use MAFFT to obtain an MSA separately for each viral sequence corpus (the same corpuses used to train our language models). We then compute the mutational frequency independently at each residue position, with respect to the wildtype sequence. To ensure high quality sequence alignments, we also restrict this computation to aligned sequences with a limited number of gap characters relative to the wildtype (0 gap characters for the influenza and coronavirus proteins and 15 gap characters for HIV Env). The mutational frequency for each amino acid at each residue was used as the measure of viral fitness for escape acquisition (acquiring mutants with higher observed frequency).

For our second fitness model, we use the EVcouplings framework (Hopf et al., 2017, 2019) (https://github.com/debbiemarkslab/EVcouplings), which leverages HMMER software for sequence alignment (Eddy, 2008) (http://hmmer.org/), to estimate the predicted fitness using both the independent and epistatic models. We train the EVcouplings models using the same sequence corpuses that we used for training our language models. We acquired mutants based on higher predicted fitness scores obtained from the independent or the epistatic model. The epistatic model incorporates pairwise residue information by learning a probabilistic model in which each residue position corresponds to a random variable over an amino acid alphabet and pairwise information potentials can encode epistatic relationships.

We use a number of pretrained protein sequence embedding models to assess the ability for generic protein embedding models to capture antigenic information. We use the pretrained soft symmetric alignment model with multitask structural training information from Bepler and Berger (Bepler and Berger, 2019) (https://github.com/tbepler/protein-sequence-embedding-iclr2019). We also use the TAPE pretrained transformer model (Rao et al., 2019) and the UniRep pretrained model (Alley et al., 2019), both from the TAPE repository (https://github.com/songlab-cal/tape). Using each pretrained model, we computed an embedding for the wildtype sequence and for each single-residue mutant. We used the *ℓ*_1_ distance between the wildtype and mutant embeddings as the semantic change score. We acquired mutants favoring higher semantic change.

### Protein structure preprocessing and visualization

We calculated the escape potential at each position within a given viral sequence by summing the value of the CSCS rank-based acquisition function, i.e., 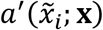, across all amino acids. We then mapped these scores from our protein sequences of interest (used in the escape prediction validation experiments) to three-dimensional structural loci. As in Doud et al. (Doud et al., 2018), we mapped the positions from WSN33 to the structure of HA H1 A/Puerto Rico/8/1934 (PDB: 1RVX) (Gamblin et al., 2004). As in Lee et al. (Lee et al., 2019), we mapped the positions from Perth09 to the structure of HA H3 A/Victoria/361/2011 (PDB: 4O5N) (Lee et al., 2014). As in Dingens et al. (Dingens et al., 2019), we used the structure of BG505 SOSIP (PDB: 5FYL) (Stewart-Jones et al., 2016). We used the structure of the closed-state Spike ectodomain (PDB: 6VXX) (Walls et al., 2020). Escape potential across each of the structures was colored using a custom generated PyMOL script and visualized with PyMOL version 2.3.3 (https://pymol.org/2/).

### Protein structure regional enrichment and depletion quantification

We quantified the enrichment or depletion of escape prediction scores within a given region of a protein sequence. We define a region as a (potentially non-contiguous) set of positions; regions of each viral protein that we considered are provided in **table S3**. Head and stalk regions for HA were determined based on the coordinates used by Kirkpatrick et al. (Kirkpatrick et al., 2018). Region positions for Env were determined using the annotation provided by UniProt (ID: QN0S5) and hypervariable loops were determined as defined by the HIV LANL database (https://www.hiv.lanl.gov/content/sequence/VAR_REG_CHAR/variable_region_characterization_explanation.html). Region positions for SARS-CoV-2 were determined using the annotation provided by UniProt (ID: P0DTC2).

To assess statistical significance, we construct a null distribution using a permutation-based procedure. We “permute” the labels corresponding to the region of interest by randomly selecting (without replacement) a set of positions that has an equal size as the region of interest; we then compute the average escape potential over this randomly selected set of positions, repeating for 100,000 permutations. We compute a *P* value by determining the average escape potential for the true region of interest and comparing it to the null distribution, where we test for both enrichment or depletion of escape potential. Enrichment or depletion is considered statistically significant if its Bonferroni-corrected *P* value is less than 0.05.

### Computational resources and hardware

Models were trained and evaluated with tensorflow 2.2.0 and Python 3.7 on Ubuntu 18.04, with access to a Nvidia Tesla V100 PCIe GPU (32GB RAM) and an Intel Xeon Gold 6130 CPU (2.10GHz, 768GB of RAM). Using CUDA-based GPU acceleration, training on the influenza HA corpus required approximately 72 hours (all times are wall time) and evaluating all possible single-residue mutant sequences for a single strain required approximately 35 minutes. Training on the HIV Env corpus required approximately 80 hours and evaluating all possible single-residue mutant sequences required approximately 90 minutes. Training on the coronavirus spike corpus required approximately 20 hours and evaluating all possible single-residue mutant sequences required approximately 10 hours.

---

**Figure S1:**
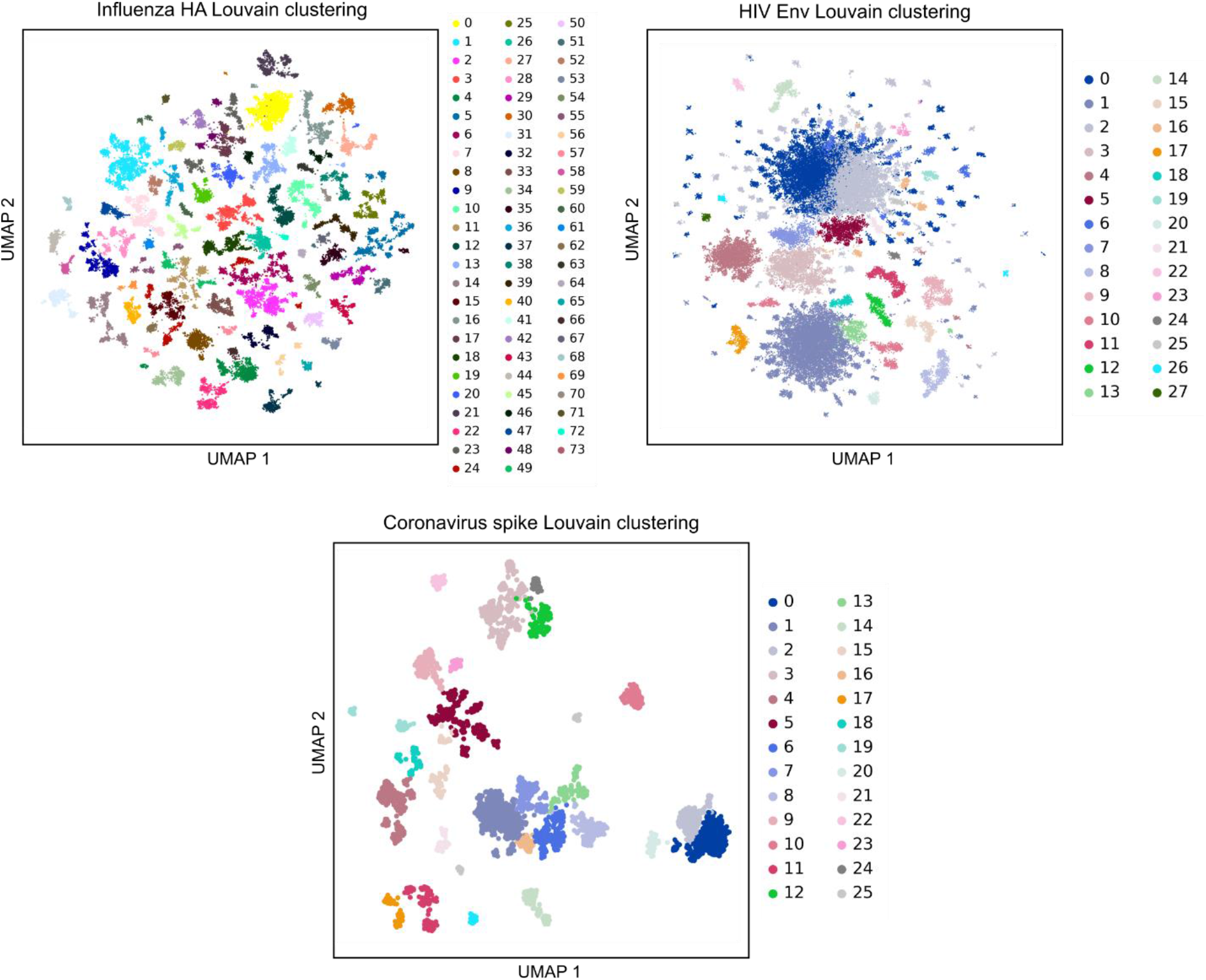
Visualization of semantic landscape Louvain clustering. Louvain cluster labels, used to evaluate cluster purity of HA subtype, HA host species, and HIV subtype, are visualized with the same UMAP coordinates as in **Figure 2**. Part of HA cluster 30 was highlighted in **Figure 2C**. Coronavirus Louvain clusters 0 and 2 were highlighted in **Figure 2G**.

**Figure S2:**
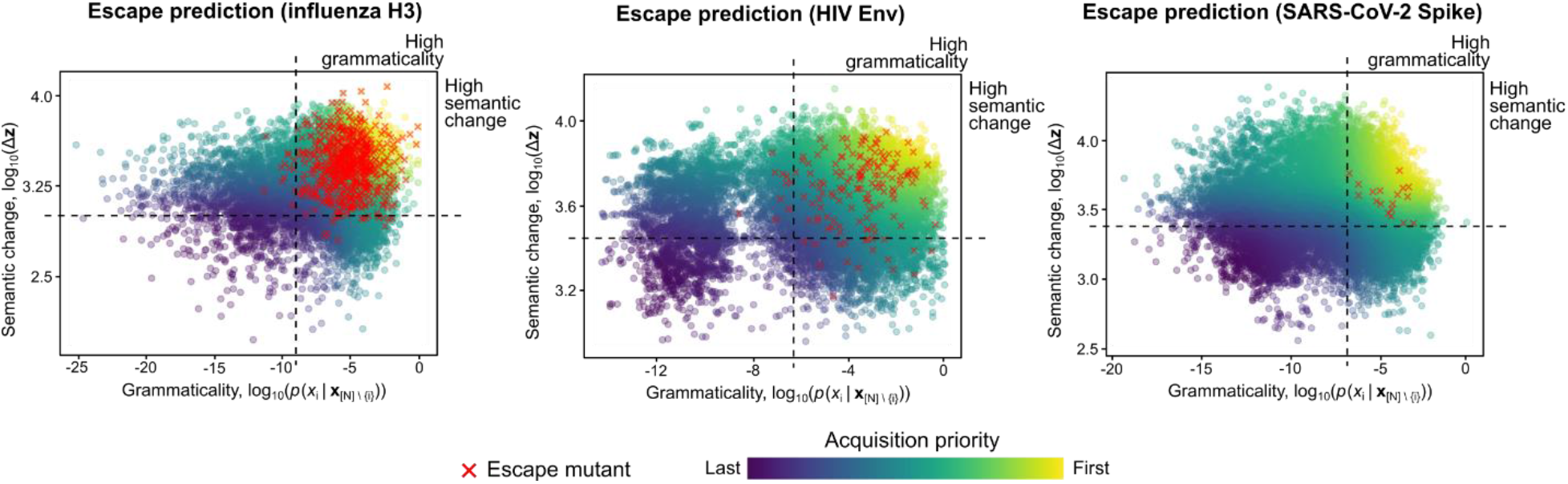
Escape mutations occupy regions of both high semantic change and grammaticality. Each point in the scatter plot corresponds to a single-residue mutation of the indicated viral protein. Points are colored by CSCS acquisition priority (**Methods**) and a red X is additionally drawn over the points that correspond to escape mutations.

**Figure S3:**
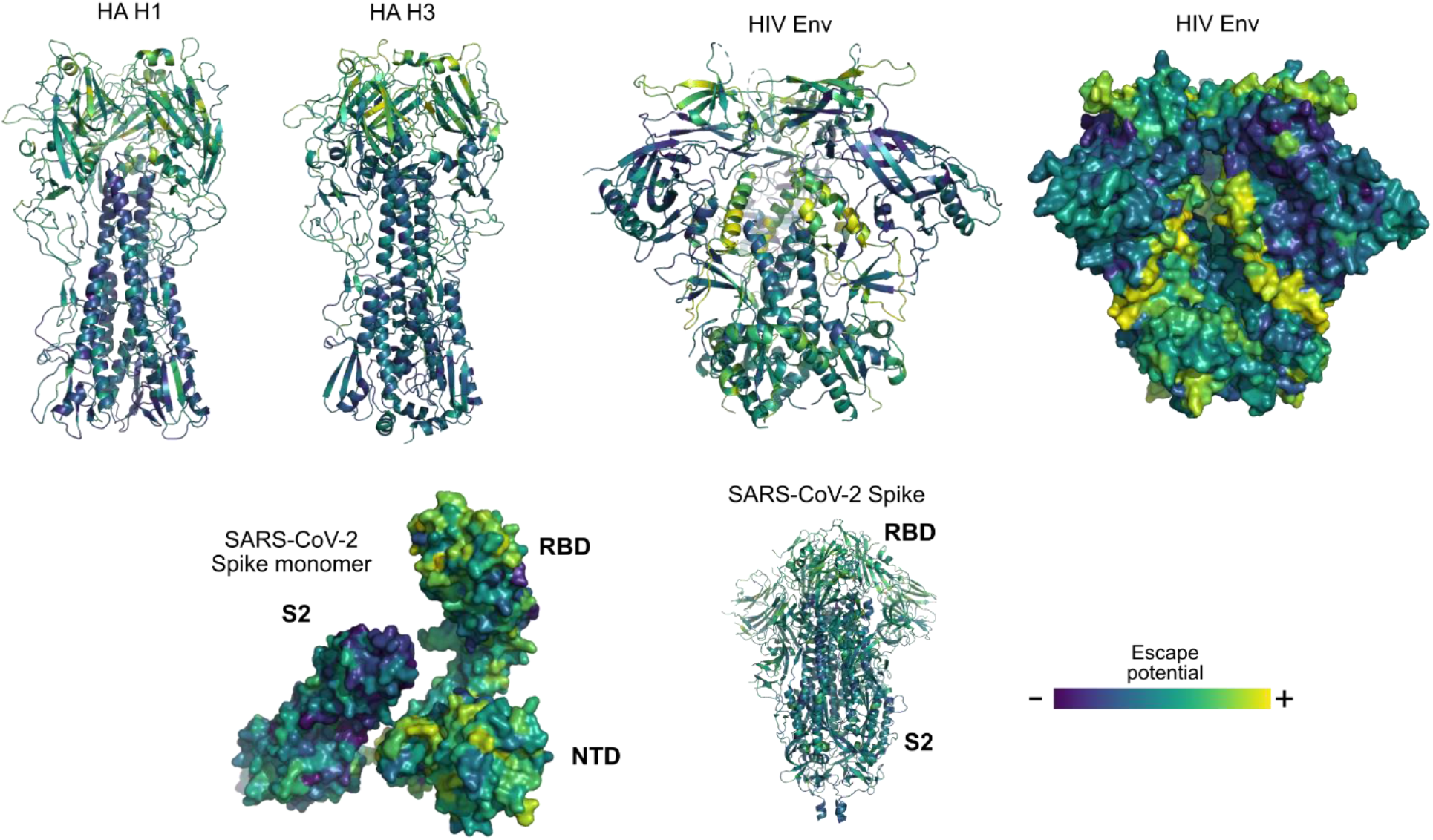
Additional protein structure visualizations. Cartoon illustration of HA H1 and HA H3; view of HIV Env as cartoon and surface oriented to illustrate the semantically important inner domain; and views of SARS-CoV-2 Spike in monomeric (surface) and trimeric form (cartoon) illustrating S2 escape depletion.

**Table S1:** Fitness correlation and *P* values. Values indicate Spearman correlation and corresponding *P* values between fitness and either semantic change or grammaticality. A *P* value of <1E-308 indicates a value that was below the floating-point precision of our computer.

| **Model** | **WSN33** | **Bei89** | **Bk79** | **Bris07L194** | **HK68** | **Mos99** | **NDako16** | **BF520** | **BG505** | **Spike** |
| --- | --- | --- | --- | --- | --- | --- | --- | --- | --- | --- |
| Semantic change<br>(Spearman r) | -0.1175 | -0.1653 | -0.4040 | -0.0051 | -0.2549 | -0.2097 | -0.0711 | -0.1024 | -0.1101 | -0.4421 |
| Grammaticality<br>(Spearman r) | 0.2789 | 0.1876 | 0.4274 | 0.4021 | 0.5396 | 0.2854 | 0.3508 | 0.2063 | 0.2684 | 0.4852 |
| Semantic change<br>(*P* value) | 2.94E-34 | 6.69E-05 | 5.02E-24 | 0.9034 | 5.38E-10 | 3.80E-07 | 0.08842 | 1.20E-30 | 1.16E-35 | <1E-308 |
| Grammaticality<br>(*P* value) | 1.08E-190 | 5.81E-06 | 5.55E-27 | 8.47E-24 | 7.97E-45 | 2.96E-12 | 4.02E-18 | 6.54E-121 | 4.85E-209 | <1E-308 |

**Table S2:** Escape prediction normalized AUC values. Normalized AUC values for escape prediction as plotted in **Figure 3B**, as well as separate AUCs for grammaticality and semantic change alone (this information is combined, as described in the **Methods** section, for our model’s full CSCS acquisition). Rows involving our model are highlighted in blue.

| **Model** | **HA H1** | **HA H3** | **Env BG505** | **Spike** |
| --- | --- | --- | --- | --- |
| MAFFT | 0.697 | 0.598 | 0.523 | 0.571 |
| EVcouplings (ind.) | 0.706 | 0.691 | 0.536 | 0.674 |
| EVcouplings (epi.) | 0.726 | 0.687 | 0.552 | 0.696 |
| Grammaticality<br>(our model) | 0.820 | 0.684 | 0.667 | 0.826 |
| Bepler | 0.660 | 0.644 | 0.561 | 0.513 |
| TAPE transformer | 0.584 | 0.526 | 0.574 | 0.634 |
| UniRep | 0.482 | 0.452 | 0.534 | 0.740 |
| Semantic change<br>(our model) | 0.664 | 0.709 | 0.622 | 0.653 |
| CSCS<br>(our model) | **0.834** | **0.771** | **0.692** | **0.856** |

**Table S3:** Escape potential regions of interest. Residue positions corresponding to the regions of interest tested for enrichment or depletion of escape potential in **Figure 4**. All ranges are inclusive.

| **Virus** | **Region name** | **Region positions** |
| --- | --- | --- |
| H1 WSN33 | Head | 59 – 291 |
|  | Stalk | 18 – 58, 292 – 528 |
| H3 Perth09 | Head | 68 – 293 |
|  | Stalk | 17 – 67, 294 – 530 |
| Env BG505 | V1 loop | 130 – 148 |
|  | V2 loop | 149 – 195 |
|  | V3 loop | 295 – 329 |
|  | V4 loop | 383 – 415 |
|  | V5 loop | 458 – 468 |
|  | CD4 binding loop | 362 – 372 |
|  | Fusion peptide | 509 – 529 |
|  | Immunosuppression | 571 – 589 |
|  | MPER | 659 – 680 |
|  | gp120 A | 33 – 139 |
|  | gp120 B | 144 – 508 |
|  | gp41 | 527 – 716 |
|  | Glycosylation | 87, 132, 136, 147, 151, 196, 233, 261, 275, 294, 300, 337, 353, 361, 384, 390, 445, 608, 615, 634 |
| Spike | S1 NTD | 13 – 303 |
|  | RBD | 319 – 541 |
|  | Fusion peptide | 788 – 806 |
|  | HR1 | 920 – 970 |
|  | HR2 | 1163 – 1202 |
|  | S2 | 686 – 1273 |
|  | Glycosylation | 17, 61, 74, 122, 149, 165, 234, 282, 331, 343, 603, 616, 657, 709, 717, 801, 1074, 1098, 1134, 1158, 1173, 1194 |

## Notes

https://github.com/brianhie/viral-mutation

### Competing Interest Statement

The authors have declared no competing interest.

